# Hypocretin underlies the evolution of sleep loss in the Mexican cavefish

**DOI:** 10.1101/122903

**Authors:** James B. Jaggard, Bethany A. Stahl, Evan Lloyd, David A. Prober, Erik R. Duboue, Alex C. Keene

**Affiliations:** Department of Biological Sciences, Florida Atlantic University, Jupiter, FL 33458; Division of Biology and Biological Engineering, California Institute of Technology, Pasadena, CA 91125, USA; Department of Embryology, Carnegie Institution for Science, Baltimore, MD, 21218; Harriet L. Wilkes Honors College, Florida Atlantic University, Jupiter, FL 33458

## Abstract

The duration of sleep varies dramatically between species, yet little is known about genetic bases or evolutionary factors driving this variation in behavior. The Mexican cavefish, *Astyanax mexicanus*, exists as surface populations that inhabit rivers, and multiple independently derived cave populations with convergent evolution on sleep loss. The number of Hypocretin/Orexin (HCRT)-positive hypothalamic neurons is increased significantly in cavefish, and HCRT is upregulated at both the transcript and protein levels. Pharmacological or genetic inhibition of HCRT signaling increases sleep duration in cavefish without affecting sleep in surface fish, suggesting enhanced HCRT signaling underlies sleep loss in cavefish. Ablation of the lateral line or starvation, manipulations that selectively promote sleep in cavefish, inhibit *hcrt* expression in cavefish while having little effect in surface fish. These findings provide the first evidence of genetic and neuronal changes that contribute to the evolution of sleep loss, and support a conserved role for HCRT in sleep regulation.

## Introduction

Sleep behavior is nearly ubiquitous throughout the animal kingdom and vital for many aspects of biological function (Campbell and Tobler, 1984; Hartmann, 1973; Musiek et al., 2015). While animals display remarkable diversity in sleep duration and architecture, little is known about the functional and evolutionary principles underlying these differences (Allada and Siegel, 2008; Capellini et al., 2008; Siegel, 2005). We previously described convergent evolution of sleep loss in the blind Mexican cavefish, *Astyanax mexicanus.* Relative to extant conspecifics that inhabit caves, independently derived cave-dwelling populations display a striking 80% reduction in total sleep with no adverse impacts on health or development (Duboué et al., 2011). The robust differences in sleep between surface and cave populations provide a unique model for investigating the genetic basis for sleep variation and identification of novel mechanisms underlying the evolution of sleep regulation.

*Astyanax mexicanus* consists of eyed surface populations that inhabit rivers in the Sierra del Abra region of Northeast Mexico, and at least 29 distinct populations of cavefish (Mitchell et al., 1977). Cavefish are derived from surface ancestors, which arose from colonization events within the past 2-5 million years (Gross, 2012; Jeffery, 2009; Keene et al., 2015). Independently evolved cave populations of *A. mexicanus* share multiple morphological and developmental phenotypes including smaller or completely absent eyes, and loss of pigmentation (Borowsky, 2008a; Gross and Wilkens, 2013; Protas et al., 2006). In addition, cavefish display an array of behavioral changes including reduced schooling, enhanced vibration attraction behavior, hyperphagia, and sleep loss (Aspiras et al., 2015; Duboué et al., 2011; Kowalko et al., 2013; Yoshizawa et al., 2010). Convergent evolution of shared traits in independent cavefish populations, combined with robust phenotypic differences with extant surface fish populations, provides a system to examine how naturally occurring variation and evolution shape complex biological traits.

While the ecological factors underlying phenotypic changes in cave populations are unclear, food availability and foraging strategy are hypothesized to be potent drivers of evolutionary change that contribute to the variation in sleep duration across animal species (Siegel, 2005). Many cave waters inhabited by *A. mexicanus* are nutrient poor compared to the above-ground rivers surrounding them (Mitchell et al., 1977), and previous field studies suggest cavefish subsist primarily off of bat guano, small insects, and organic matter washed into the cave by seasonal floods (Keene et al., 2015; Mitchell et al., 1977). Following starvation, cave derived fish have a slower rate of weight loss compared to surface conspecific, suggesting that a reduced metabolism may account, in part, for adaptation to cave life (Aspiras et al., 2015). We previously found that sleep is increased in cavefish during periods of prolonged starvation, raising the possibility that cavefish suppress sleep to forage during the wet season when food is plentiful, and increase sleep to conserve energy during the dry season when food is less abundant (Jaggard et al., 2017). Therefore, sleep loss in cavefish appears to be an evolved feature as a consequence of changes in food availability, providing a model to examine interactions between sleep and metabolism.

Despite the robust phenotypic differences in sleep between *A. mexicanus* surface and cave populations, little is known about the genetic or neuronal mechanisms underlying the evolution of sleep loss in cavefish. Many behaviors that are altered in cavefish are regulated by the hypothalamus, which is enlarged in cavefish (Menuet et al., 2007). Here, we investigate the role of Hypocretin/Orexin (HCRT), a highly conserved hypothalamic neuropeptide known to promote wakefulness. Deficiencies in HCRT signaling are associated with altered sleep and narcolepsy associated phenotypes in diverse vertebrate organisms, including mammals (Chemelli et al., 1999; Lin et al., 1999) and zebrafish (Elbaz et al., 2012; Yokogawa et al., 2007). Conversely, overexpression of Hcrt or stimulation of *hcrt*-expressing neurons promotes arousal in mammals (Adamantidis and de Lecea, 2008; Mieda et al., 2004) and zebrafish (Chen et al., 2016; Prober et al., 2006; Singh et al., 2015). Here we provide evidence that alteration of the Hcrt system contributes to sleep loss in cavefish. We find that pharmacologic or genetic disruption of HCRT signaling selectively reduces sleep in cavefish but not in their surface morphs. Further, we show that HCRT expression is down-regulated in cavefish in response to sleep-promoting manipulations including starvation and ablation of the lateral line (Jaggard et al., 2017). Together, these findings suggest that plasticity of HCRT function contributes to evolved differences in sleep regulation in Mexican cavefish.

## Results

Sleep is dramatically reduced in adult Pachón cavefish compared to surface fish counterparts (Fig. 1A, B) (Jaggard et al., 2017; Yoshizawa et al., 2015). We compared sequence homology between surface fish and cavefish by a bioinformatic analysis of the sequences from the cavefish genome (McGaugh et al., 2014) and available full-length transcriptomic sequences (Gross et al., 2013). Alignment of the HCRT neuropeptide reveals that the *Astyanax* ortholog shares high amino acid sequence similarity to other fish species (35-48% percent identity) and mammals (35% percent identity), including conservation of domains that give rise to the HCRT neuropeptides (Fig. 1-S1A; Wall and Volkoff, 2013). The HCRT peptide sequences of surface and Pachón cavefish are 100% identical. To determine if *hcrt* expression differs between adult surface fish and cavefish, we measured transcript levels in whole-brain extracts by quantitative PCR (qPCR). Expression of housekeeping genes such as *rpl3a* and *gapdh* were comparable between both forms (Fig. 1-S1B). By contrast, expression of *hcrt* was significantly elevated in Pachón cavefish to over three-fold the levels of surface fish, raising the possibility that up regulation of *hcrt* underlies sleep loss in Pachón cavefish (Fig. 1C). To determine if cavefish have higher Hcrt protein levels and/or more Hcrt-positive neurons, we performed immunolabeling of serial-sectioned brains, and examined the number of Hcrt-positive cell bodies and the relative fluorescence of each cell under fed conditions (Fig 1- S2A-D). The number of HCRT-positive cell bodies was significantly higher in Pachón cavefish compared to surface fish (Fig. 1D). Further, quantification of fluorescence intensity of individual cells revealed that Hcrt-positive cells produce higher levels of Hcrt peptide in cavefish (Fig. 1E-I). Increased levels of HCRT protein were also observed in 5 day post fertilization (dpf) larvae, suggesting the change in peptide levels were present at the time fish begin consuming food (Fig. 1-S3).

**Figure 1:**
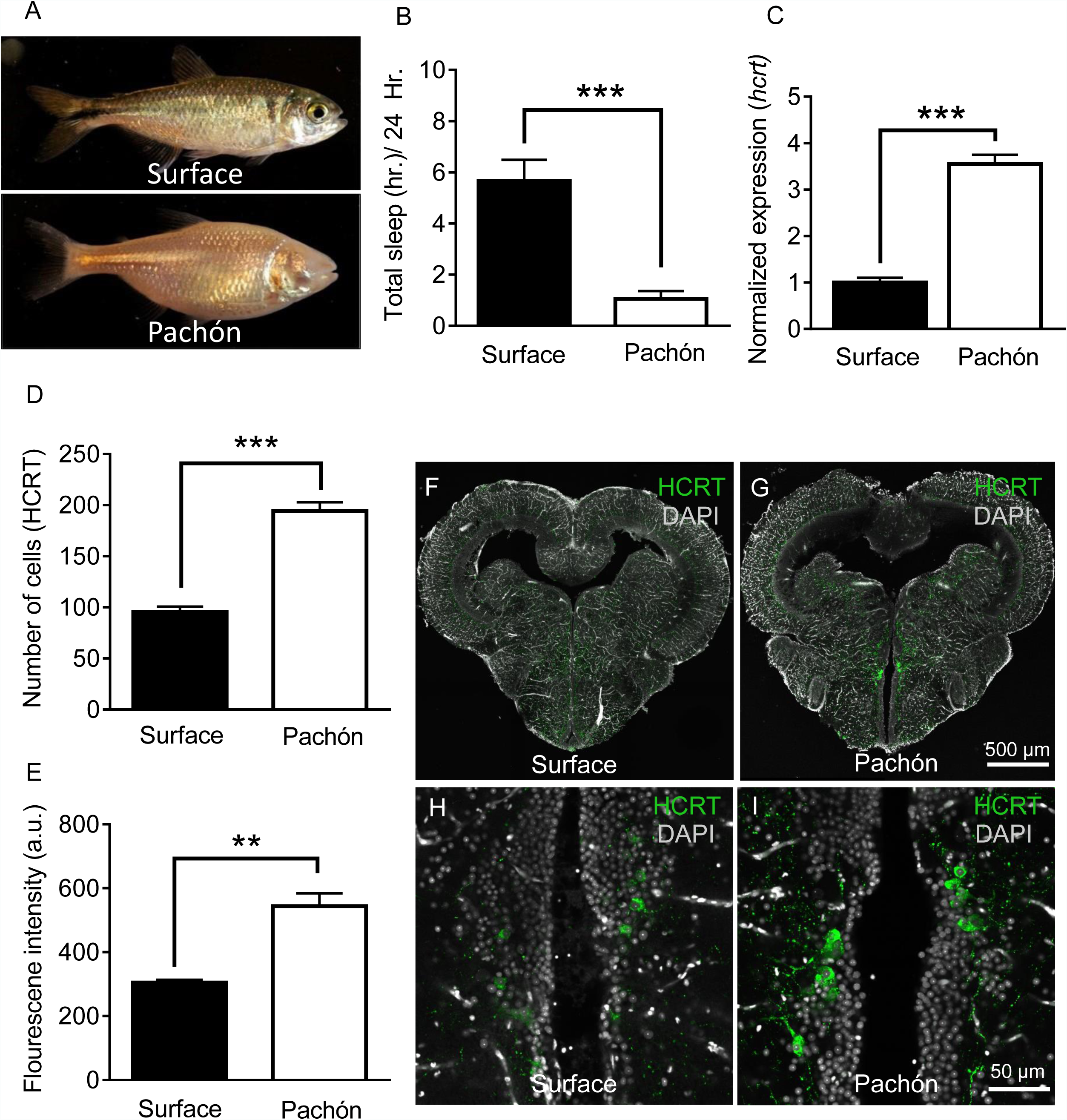
Hypocretin transcript and peptide levels are elevated in Pachón cavefish. **A.** Representative images of surface fish and Pachón cavefish **B.** Sleep duration is significantly reduced in Pachón cavefish compared to surface morph (Unpaired t-test, t=5.56, n=26, P< 0.0001). **C.** Expression of *hcrt* normalized by GAPDH and *rpl13α* in adult whole-brain extracts is significantly enhanced in Pachón cavefish compared to surface fish (Unpaired t-test, t=11.15, n=8, P< 0.0001). **D.** HCRT neuropeptide signal is significantly increased in Pachón cavefish compared to surface fish (Unpaired t-test, t=5.94, n=8, P< 0.001). E. The number of HCRT-positive cells in the hypothalamus is significantly increased in cavefish compared to surface fish (Unpaired t-test, t=9.984, n=8, P< 0.0001). **F-I.** Representative 2 µm confocal images from coronal slices of surface fish or Pachón brains immunostained with anti-HCRT (green) and DAPI (white) **F.** Surface whole brain coronal slice. **G.** Pachón whole brain coronal slice. **H.** Surface fish dorsal hypothalamus containing HCRT positive cells **I.** Pachón cavefish dorsal hypothalamus containing HCRT neurons in view. Scale bar denotes 500µm (F,G); 50µm (H,I).

To assess more directly the contribution of Hcrt signaling to sleep loss, we measured the effect of HCRT receptor inhibition on sleep in adult surface fish and Pachón cavefish. While mammals possess two Hcrt receptors (Hcrtr1 and Hcrtr2), the genomes of zebrafish (Prober et al., 2006; Yokogawa et al., 2007), as well as *A. mexicanus* cavefish and surface fish (McGaugh et al., 2014), contain only *hcrtr2*. Importantly, Hcrtr2 is thought to be evolutionarily more ancient than Hcrtr1 (McGaugh et al., 2014; Wong et al., 2011).

Fish from both populations were treated with the selective Hcrtr2 pharmacological inhibitor TCS0X229 (Kummangal et al., 2013; Plaza-Zabala et al., 2012). Surface fish sleep was unchanged in the presence of 1 µM or 10 µM TCS0X229 (Fig. 2A, C). In contrast, treatment of Pachón cavefish with TCS0X229 increased sleep duration compared to DMSO vehicle control (Fig. 2B, C). While these results do not exclude the possibility that HCRT regulates sleep in surface fish, the sleep-promoting effect of TCS0X229 in Pachón cavefish suggests these fish are more sensitive to changes in HCRT signaling than surface fish. Treatment with TCS0X229 had no effect on waking activity (the amount of locomotor activity while awake) in surface fish or cavefish, suggesting that the increased quiescence observed in cavefish after drug treatment is not due to lethargy (Fig. 2D). Further analysis revealed that sleep-promoting effects of TCS0X229 in cavefish can be attributed to both an increase in bout number and bout duration, suggesting that inhibition of Hcrt signaling affects both sleep onset and maintenance (Fig. 2E, F). Taken together, these findings support the notion that elevated HCRT signaling in cavefish underlies, in part, the evolution of sleep loss.

**Figure 2.**
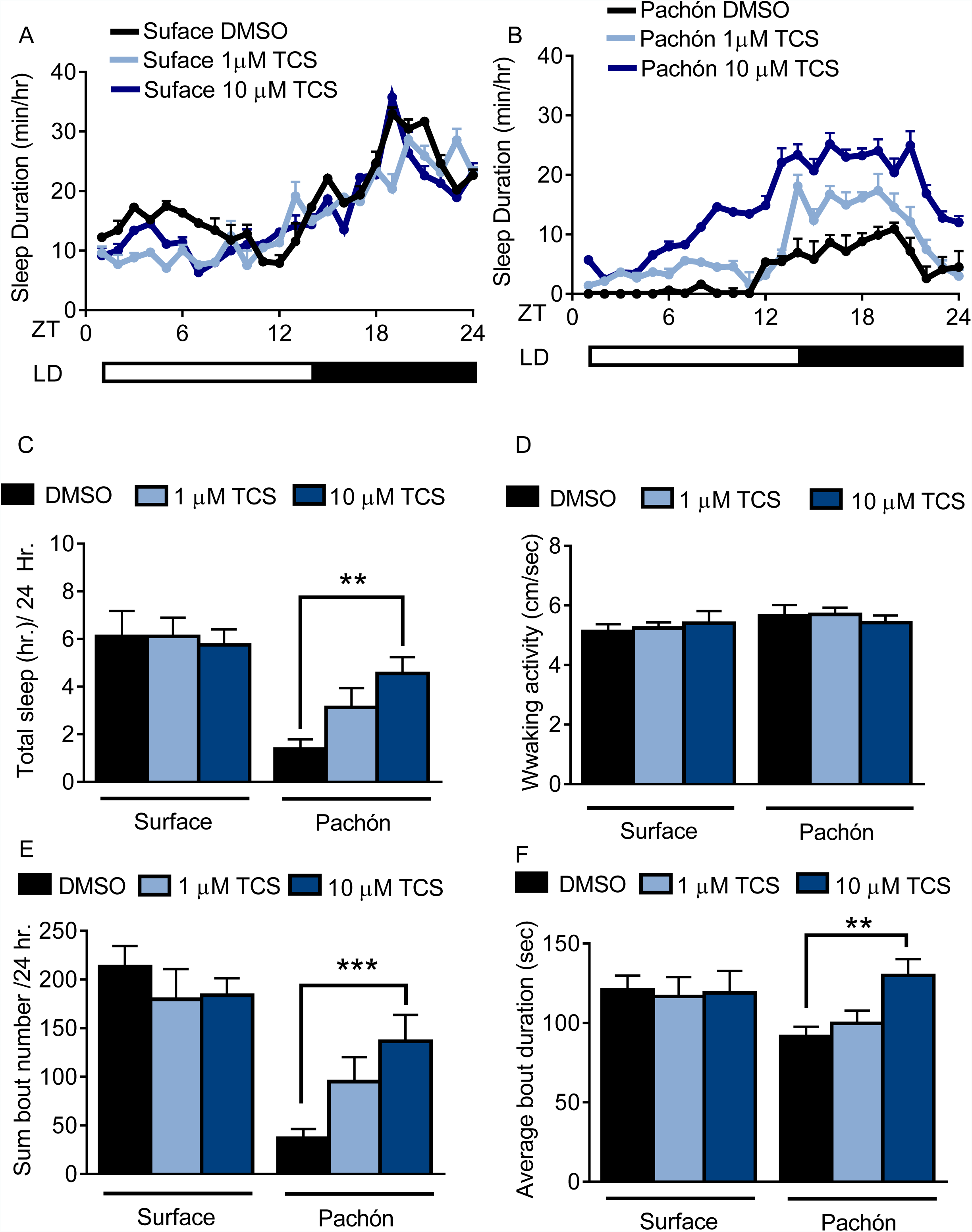
Pharmacological inhibition of HCRT Receptor 2 promotes sleep in Pachón cavefish. A,B. Twenty-four hour sleep profile in surface fish (A) Pachón cavefish (B) treated with DMSO (black), 1µm TCS (light blue) or 10µm TCS (dark blue). **C.** TCS treatment does not affect total sleep duration in surface fish (1µM p>0.999, n=12, 10 uM P>0.941, n=12). Pachón cavefish teated with 1µM TCS trended towards increased sleep (P>0.178, n=12) while treatment with 10µM significantly increased sleep (P< 0.01, n=13; F(1, 73) = 25.00,) compared to control treated fish. **D.** Waking activity was not significantly altered in surface fish or cavefish or in response to drug treatment, 2-way ANOVA, (F(1, 73) = 2.73, P>0.103, n=79) **E.** Treatment with TCS did not affect average sleep bout duration in surface fish (1 µM TCS, P>0.430, n=12; 10µM TCS, P>0.518, n=12) Treatment of Pachón cavefish with 1µM TCS trended towards increased bout duration, P>0.051, n=12, while 10µM TCS treatment significantly increased bout duration in Pachón cavefish, (P< 0.01, n=13; F(1, 73) = 47.42). **F.** TCS treatment did not affect total sleep bout number in surface fish 1µM TCS, P>0.976, n=12; 10µM TCS, P>0.998, n=12). In Pachón cavefish, treatment with 1µM TCS did not affect sleep bout number (P>0.828, n=12). Treatment with 10µM TCS significantly increased bout duration in Pachón cavefish,(P< 0.001, n=13; 2-way ANOVA, F(1, 68) = 3.309).

Sleep loss in *A. mexicanus* cavefish populations is found at all developmental stages, from larval and juvenile forms to adults (Duboué et al., 2011; Yoshizawa, 2015). The small size of young fry (25 dpf) and ability to perform higher throughput analysis make them an excellent model for investigating the effects of drugs on sleep. Previous drug screens have been carried out using larval and juvenile zebrafish and *A. mexicanus* using standard concentrations between 1-30 µM for all drugs (Duboué et al., 2011; Rihel et al., 2010). We therefore selected additional pharmacological modulators of HCRTR2 based on permeability to blood-brain barrier and affinity for HCRTR for testing in fry (Fig. 3-S1A). Fish from both populations were bathed in the selective HCRTR2 pharmacological inhibitors TCS0X229 (Kummangal et al., 2013; Plaza-Zabala et al., 2012) or *N*-Ethyl-2-[(6-methoxy-3-pyridinyl)[(2-methylphenyl)sulfonyl]amino]-*N*-(3- pyridinylmethyl)-acetamide (EMPA) (Malherbe et al., 2009; Mochizuki et al., 2011), or the HCRTR1/2 antagonist Suvorexant (Betschart et al., 2013; Hoyer et al., 2013) (Fig 3A). A dose-response assay was carried out in juvenile fish for TCSOX229 (Fig. 3-S1B-D) and found a significant effect in Pachón cavefish at concentrations ranging from 10-30 µM. Therefore, all three antagonists were tested for their effect on sleep in surface and cavefish at a dose of 30 µM. None of the three antagonists affected sleep in surface fish, whereas they all significantly increased sleep in cavefish (Fig 3B). Waking activity was not affected by treatment with any of the antagonists (Fig 3C), suggesting that the increased sleep in cavefish is not due increased lethargy. Further, all three HCRT antagonists significantly increased sleep bout number and duration (Fig. 3D, E). Therefore, larval and adult Pachón cavefish are more sensitive to changes in HCRT signaling than surface fish.

**Figure 3:**
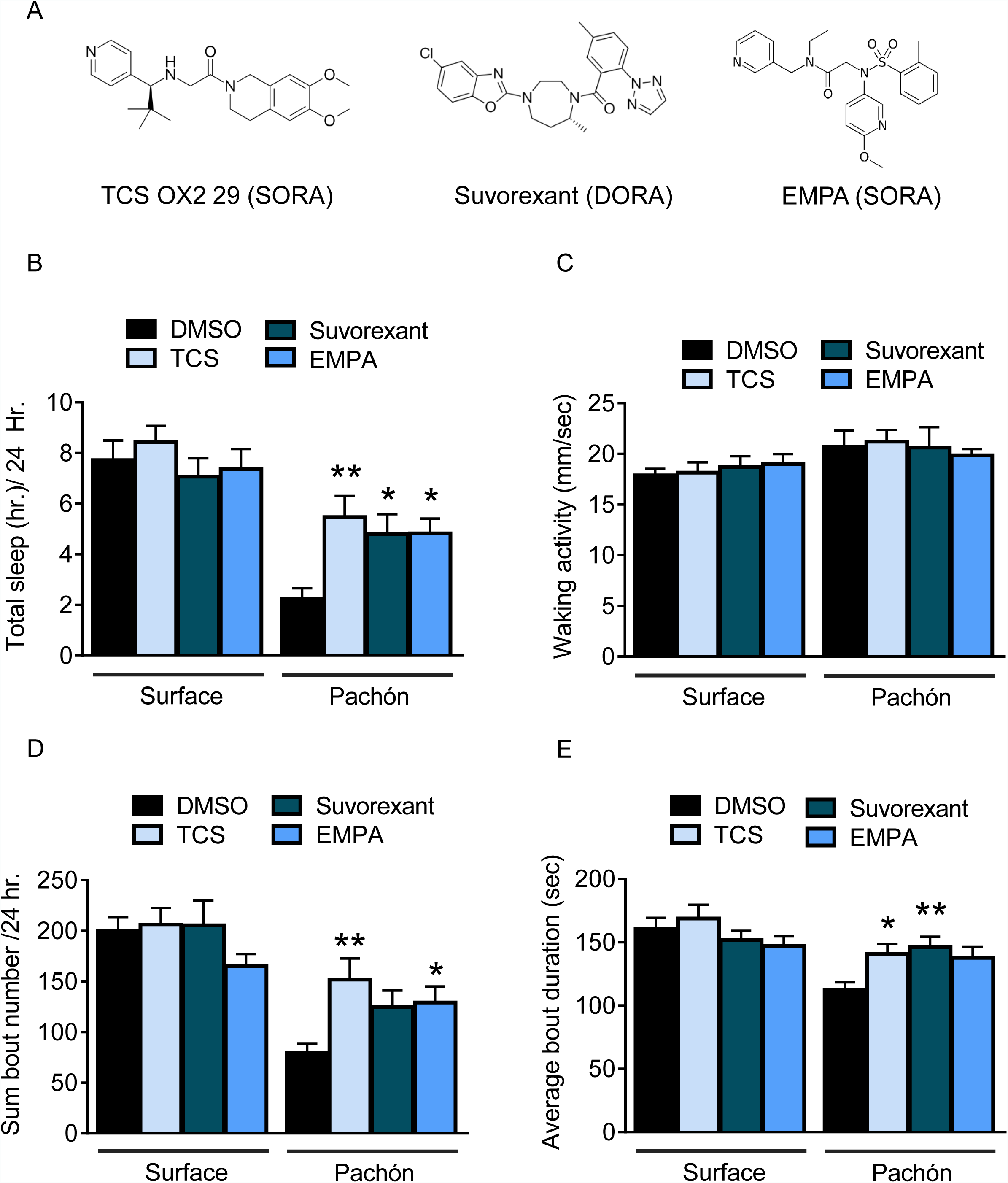
Panel of pharmacological HCRT antagonists reveals wake-promoting role of HCRT in juvenile cavefish A. Two SORAs and one DORA were used to inhibit binding of HCRT to its receptor. **B.** Total sleep is not significantly altered in surface fish with the treatment of HCRT antagonists (TCS, P=0.814, n=18; Suvorexant, P=0.865, n=18; EMPA, P=0.972, n=19). Pachón cavefish significantly increased total sleep in response to HCRT antagonists (TCS, P=0.004, n=18; Suvorexant, P=0.032, n=19, EMPA, P=0.027, n=18; 2-way ANOVA, F(1, 135)=43.70, P< 0.0001). **B.** Waking activity was not significantly altered in any fish or in response to drug treatment, Surface TCS, P=0.996, n=18; Suvorexant, P=0.925, n=18; EMPA, P=0.816, n=19; Pachón, TCS, P=0.980, n=18; Suvorexant, P>0.999, n=19; EMPA, P=0.917, n=18; 2-way ANOVA, (F_(1, 135)_ = 7.21, P=0.008) **C.** The total number of sleep bouts over 24 hours of drug treatment did not significantly change in Surface fish (TCS, P=0.814, n=18; Suvorexant, P=0.865, n=18; EMPA, P=0.972, n=19). Total sleep bouts were significantly increased in Pachón cavefish with TCS, P=0.002, n=18; and EMPA, P=0.041, n=18, Suvorexant trended towards significance, P=0.071, n=19 (2-way ANOVA, F(1, 135)=45.27, P< 0.0001). **D.** Average sleep bout length was not significantly different in Surface fish treated with HCRT antagonists (TCS, P=0.822, n=18; Suvorexant, P=0.805, n=18; EMPA, P=0.236, n=19). Pachón cavefish significantly increased their average sleep bout lengths with TCS, P=0.039, N=18; and with Suvorexant, P=0.009, n=19. EMPA trended towards significance, P=0.078, n=18 (2-way ANOVA, F(1, 135)=14.06, P=0.0003).

To determine whether enhanced HCRTR2 signaling was sufficient to induce sleep loss, we bathed 25 dpf fry in the HCRTR2 selective agonist YNT-185 (Fig 3-S1). While the effect of this drug on sleep has not been previously tested in fish, it promotes wakefulness in mice (Irukayama-Tomobe et al., 2017; Nagahara et al., 2015). Treatment with YNT-185 significantly reduced sleep in surface fish, without affecting sleep in cavefish where HCRT levels are naturally elevated (Fig 2-S1). Activity during waking bouts was not significantly altered with treatment of YNT-185 in either surface or Pachón cavefish, suggesting that the reduction in total sleep of surface fish was not due to hyperactivity. Further, YNT-185 treatment significantly reduced the total number of sleep bouts (Fig 3-S1). Taken together, these findings support the notion that elevated HCRT signaling in larval and adult cavefish underlies the evolution of sleep loss.

As an alternative approach to test the hypothesis that sleep loss in cavefish is due to increased Hcrt signaling, we selectively knocked-down HCRT using morpholino antisense oligonucleotides (MOs) and measured the effect on sleep in both surface and cave fish. MOs have been effectively used in zebrafish and *A. mexicnaus* to knock-down gene function (Bill et al., 2009). The effect of MO injection on gene expression is typically limited to ˜four days post injection, and we first verified that sleep differences are present at this early stage. We found that at four dpf, sleep in Pachòn cavefish is significantly reduced compared to age-matched surface fish (Fig 4A). Injection of 2 ng/embryo HCRT MO increased sleep in cavefish compared to fish injected with scrambled MO control, whereas knock-down of Hcrt using the same MO had no observable effect on surface fish (Fig. 4A). The mortality of fish injected with 2 ng HCRT MO, 2 ng scrambled MO, and non-injected controls did not differ in either Surface fish (43% in Wild type, 39% scramble MO, and 37% survival at 96 hpf) or in Pachón cavefish (36% tWild type, 33% scramble MO, and 29% HCRT MO) indicating at the concentration used, there is no generalized effect of injection procedure or MO treatment on survival. Morpholino treatment did not affect activity during wake bouts in surface fish or Pachón cavefish (Fig. 4B). Analysis of sleep architecture revealed that injection of 2 ng HCRT MO increased total sleep bout number and sleep bout duration in Pachón cavefish, though not to levels of surface fish. Therefore, these findings support the notion that elevated levels of HCRT promote sleep in Pachón cavefish.

**Figure 4:**
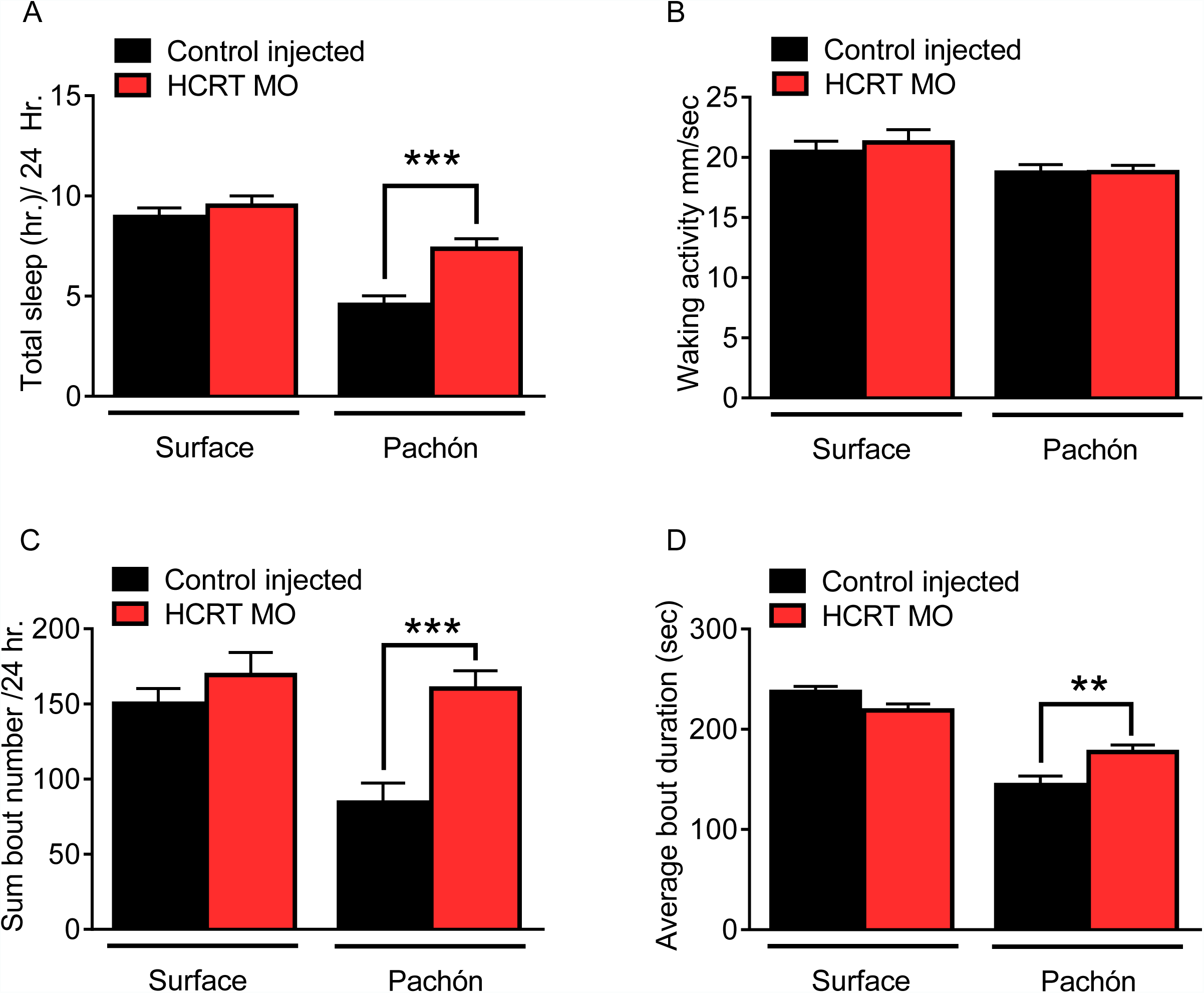
Transient knockdown of HCRT increases sleep in 4 dpf cavefish A. Morpholino knock down of HCRT does not alter total sleep in Surface fish, P=0.640, n=34, while in cavefish total sleep time was significantly increased with HCRT knockdown, P=0.0002, n=40; 2-way ANOVA, F(1,133)=45.82, P< 0.0001 **B.** Waking activity was not significantly altered in either Surface fish (P=0.343, n=34) or Pachón cavefish (P=0.084, n=40; 2-way ANOVA, F(1, 133)=5.807). **C.** Total number of sleep bouts in Surface fish was not significantly different from injected controls, P=0.459, n=34. While in Pachón cavefish, total sleep bouts over 24 hours was significantly increased in HCRT MO fish compared to control fish (P=0.0004, n=40; 2-way ANOVA, F(1,133)=8.295, P=0.004). **D.** Average sleep bout duration was not different between controls and HCRT MO injected Surface fish (P=0.081, n=34). There was a significant increase in average bout duration In Pachón HCRT MO injected fish compared to their respective controls (P=0.004, n=40; 2-way ANOVA, F(1,133)=13.61, P=0.0003).

To further validate a role for HCRT in sleep regulation we sought to genetically inhibit the activity HCRT neurons and measure the effect on sleep. The GAL4/UAS system has been widely in *Drosophila* and zebrafish to manipulate gene expression with spatial specificity (Asakawa and Kawakami, 2008; Brand and Perrimon, 1993; Scheer and Campos-Ortega, 1999). In zebrafish, co-injection of separate plasmids containing GAL4 and UAS transgenes flanked with Tol2 transposon sequences (Kawakami et al., 2000). To inhibit the activity of HCRT neurons, we co-injected embryos with plasmids in which a 1kb fragment of the zebrafish *hcrt* promoter drives expression of Gal4 (*hcrt:Gal4*) and a second in which UAS elements regulate expression of Botulinum toxin (Botx), which inhibits neurotransmission by cleaving SNARE proteins required for synaptic release (Auer et al., 2015; Brunger et al., 2008; Levitas-Djerbi et al., 2015). Embryos were injected, raised under standard conditions, and then tested for sleep at 25 dpf. Following sleep measurements, brains of individual fish were dissected and the number GFP-expressing neurons were quantified. No expression was observed in fish injected with UAS-BoTxBLC-GFP alone, indicating that BoTxBLC-GFP is not expressed in the absence of *hcrt:Gal4* (Fig 5A,C). Further, all GFP-positive neurons were co-labeled by a Hcrt-specific antibody, demonstrating that this approach specifically targets BoTxBLC transgene expression to HCRT neurons (Fig 5B,D). The total sleep in fish expressing hcrt:GAL4; UAS-BoTxTxBLC- GFP were compared to wild type fish or fish injected with UAS-BoTxTxBLC-GFP alone. Sleep was significantly increased in experimental Pachón cavefish (hcrt:GAL4:UAS-BoTxTxBLC-GFP) compared to both control groups, while there was no effect in surface fish (Fig 5E). Waking activity in surface fish and cavefish was not changed between both control groups or fish expressing hcrt:GAL4; UAS-BoTxBLC-GFP, suggesting that the increased quiescence from neuronal silencing is not due to lethargy (Fig. 5F).

**Figure 5:**
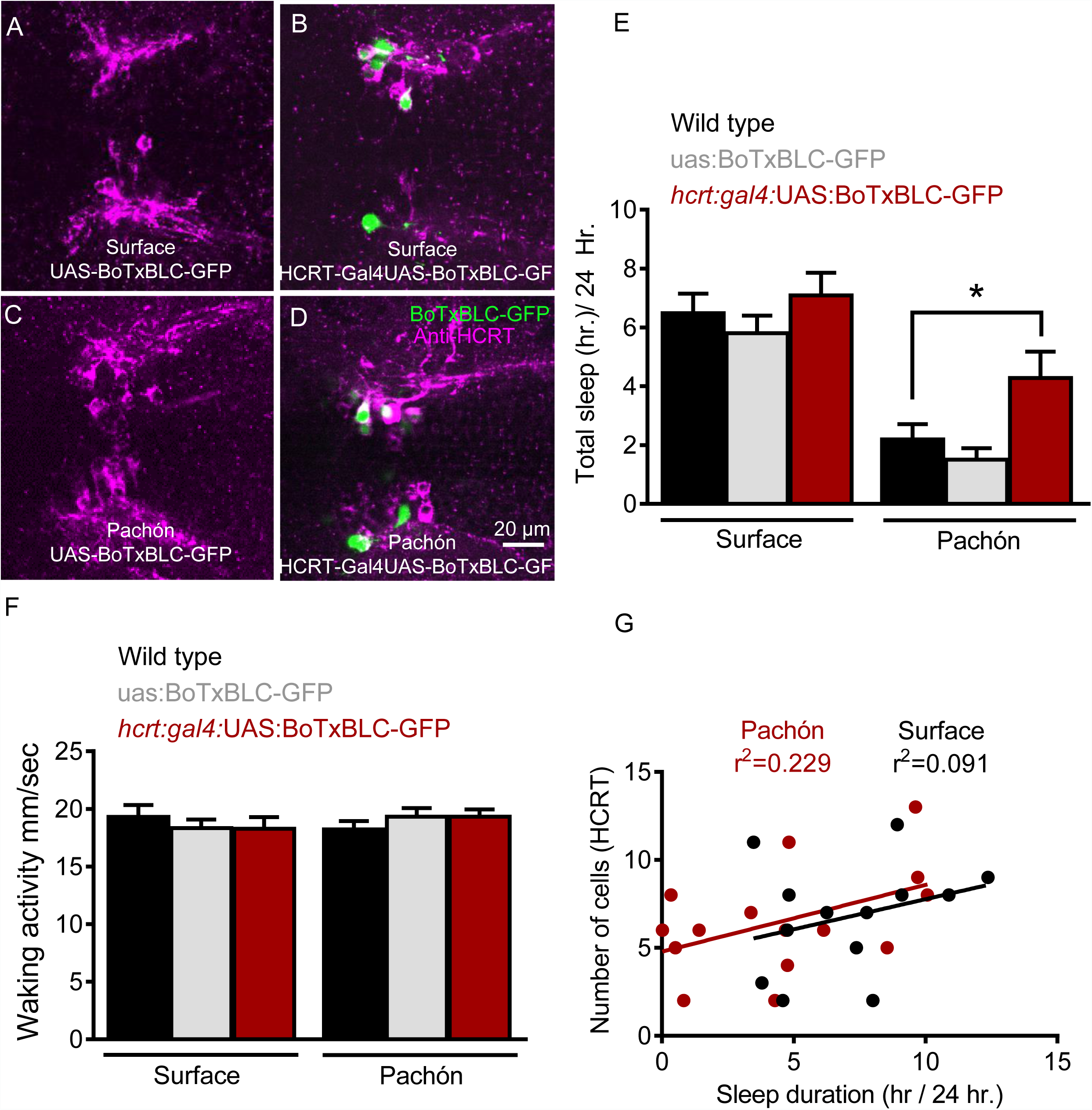
Genetic silencing of subsets of HCRT neurons selectively increases sleep in Pachón cavefish A. Surface UAS-BoTx-BLC-GFP **B.** Surface hcrt:GAL4-UAS-BoTx-BLC-GFP **C.** Pachón UAS-BoTx-BLC-GFP **D.** Pachón hcrt:GAL4-UAS-BoTx-BLC-GFP **E.** Neuronal silencing of HCRT cells did not alter total sleep duration in Surface fish (Wild Type*UAS-BoTx- BLC-GFP, P=0.989, n=23; UAS-BoTx-BLC-GFP* hcrt:GAL4-UAS-BoTx-BLC-GFP, P=0.713, n=14). Pachón cavefish increased their total sleep when HCRT cells were silenced (Wild Type*UAS-BoTx-BLC-GFP, P=0.667, n=21; UAS-BoTx-BLC-GFP* hcrt:GAL4-UAS-BoTx-BLC- GFP, P=0.0426, n=15; 2-way ANOVA, F(1,91)=65.68, P< 0.0001). **F.** Waking activity was not altered in either Surface or Pachón cavefish with neuronal silencing of HCRT (Surface fish Wild Type*UAS-BoTx-BLC-GFP, P=0.616, n=23; UAS-BoTx-BLC-GFP* hcrt:GAL4-UAS-BoTx-BLC- GFP, P=0.642, n=14; Pachón Wild Type*UAS-BoTx-BLC-GFP, P=0.587 n=21; UAS-BoTx-BLC- GFP* hcrt:GAL4-UAS-BoTx-BLC-GFP, P=0.612, n=15; 2-way ANOVA, F(1,91)=0.206, P=0.650).**G.** Regression analysis revealed there was a subtle trend of increased sleep with more HCRT neurons silenced as quantified with GFP signal (R2=0.091, P=0.314, n=14). The same regression analysis in Pachón cavefish revealed a much more robust correlation to increased sleep in more silenced HCRT cells (R2=0.229, P=0.0871, N=15).

Silencing subsets of HCRT neurons increased sleep in Pachón cavefish by increasing bout number (Fig 5A-S) without affecting bout duration (Fig 5B-S). Quantification of labeled cells reveal 10.2% of hcrt:GAL4; UAS-BoTxBLC-GFP cavefish HCRT neurons express GFP in an average of 7.5 cells per expressing animal. Fish not expressing GFP were discarded from analysis. Similarly, 11.5% of hcrt:GAL4; UAS-BoTxBLC-GFP surface fish express GFP in HCRT neurons with an average of 6.7 cells per expressing animal. Regression analysis of the number of neurons expressing BoTxBLC-GFP significantly correlated with total sleep duration (R^2^=0.229) in Pachón cavefish, consistent with the notion that increased HCRT signaling is associated with sleep loss (Fig 5G). A weak correlation between BoTx-expressing HCRT neurons and sleep duration (R^2^=0.091) was also observed in surface fish Fig 5G, consistent with the notion that HCRT may also regulates sleep in surface fish, though to a lesser extent than in cavefish populations.

Hypocretin neurons are modulated by sensory stimuli and feeding state, indicating that they are involved in the integration of environmental cues with sleep regulation (Appelbaum et al., 2007; Mileykovskiy et al., 2005). The number of mechanosensory neuromasts that comprise the lateral line, a neuromodulatory system used to detect waterflow, are increased in cavefish. This evolved trait is hypothesized to allow for an enhanced ability to forage, object detection, and social behaviors in the absence of eyes (Kowalko et al., 2013; Kulpa et al., 2015; Yoshizawa et al., 2010). We previously reported that ablation of the lateral line restores sleep to Pachón cavefish without affecting sleep in surface fish, raising the possibility that lateral input modulates HCRT signaling in cavefish to suppress sleep (Jaggard et al., 2017). To investigate the effects of lateral line input on HCRT, we pre-treated adult fish in the ototoxic antibiotic gentamicin, which effectively ablates the lateral line (Van Trump et al., 2010), and assayed sleep in adult cave and surface fish. In agreement with previous findings, gentamicin treatment fully ablated the lateral line (Fig. 6A-D) and restored sleep in cavefish without affecting sleep in surface fish (Jaggard et al., 2017). To determine the effect of lateral line ablation on HCRT regulation, we quantified *hcrt* mRNA and protein levels in adult cave and surface fish following gentamicin treatment. Quantitative PCR analysis revealed that *hcrt* expression was significantly reduced in cavefish treated with gentamicin to levels similar to untreated surface fish (Fig. 6E). We observed a non-significant decrease in *hcrt* expression following gentamicin treatment in surface fish, but this effect was much smaller than that for cavefish. Using a HCRT-specific antibody and immunohistochemistry, we found that lateral line ablation does not significantly affect the number of HCRT-positive hypothalamic neurons in surface or cavefish, but does significantly reduce the level HCRT protein within each cell in cavefish but not in surface fish (Fig. 6F-K). Together, these findings suggest that sensory input from the lateral line promotes sleep and *hcrt* expression in Pachón cavefish, providing a link between sensory input and transcriptional regulation of a wake-promoting factor.

**Figure 6:**
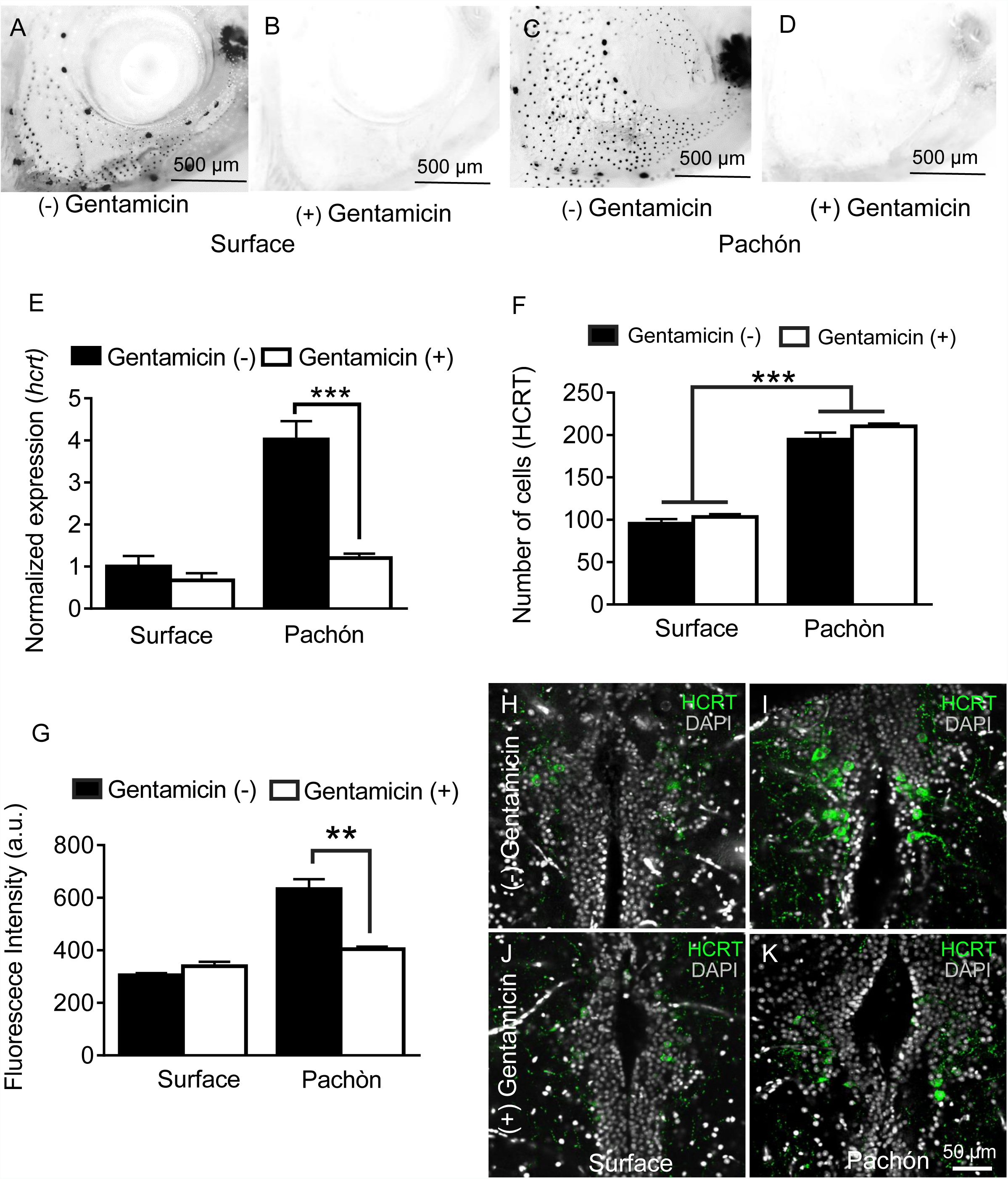
Chemical ablation of mechanosensory lateral line reduces HCRT levels in Pachón cavefish. A-D. Photomicrographs of surface fish cranial regions stained with DASPEI to reveal lateral line mechanosensory neuromasts. Treatment with gentamicin ablates lateral line neuromasts in surface fish (B) and Pachón cavefish (D) **E.** Gentamicin treatment has no significant effect on *hcrt* expression in surface fish (P>0.635, n=8) while in Pachón cavefish gentamicin treatment significantly reduces *hcrt* expression, restoring surface-like levels. (Pachón treated vs. untreated, P< 0.0001; Pachón treated vs. surface untreated, P>0.635, n=8, F(1,28)=21.28). **F.** Fluorescent intensity per hypothalamic HCRT-cell was not altered with gentamicin treatment in surface fish, P=0.590, n=4. In Pachón cavefish, HCRT neuropeptide levels are significantly lower following gentamicin treatment (P< 0.0001, n=4; 2-way ANOVA, F (1, 13) =0.0001 **G.** Gentamicin treatment has no effect on total number of HCRT cell number in either surface or Pachón cavefish (P>0.494, n=8, F (1, 13C) = 0.4967). **H-K**. Representative 2 µm confocal images of the dorsal hypothalamic region in surface fish and Pachón cavefish immunostained with HCRT (green) and DAPI (white) **H.** Surface control **I.** Pachón control **J.** Surface gentamicin K. Pachón gentamicin. Scale bar= 50 µm

In addition to its potent role in sleep regulation, *hcrt* promotes food consumption in fish and mammals (Penney and Volkoff, 2014; Tsujino and Sakurai, 2013; Yokobori et al., 2011). We previously reported that prolonged starvation increases sleep in cavefish without affecting sleep in surface fish (Jaggard et al., 2017), but the role of HCRT in feeding-state dependent modulation of sleep-wake cycles has not been investigated. Quantitative PCR analysis from whole-brain extracts revealed that *hcrt* transcript is significantly reduced in cavefish following 30 days of starvation; however, the same treatment does not affect *hcrt* transcription in surface fish, indicating that cavefish are more sensitive to starvation-dependent changes in HCRT (Fig. 7A). To determine whether HCRT neuropeptide is produced in a greater number of cells during starvation, we quantified HCRT-positive neurons in fed and starved state (Fig. 7B-G). Similar to lateral line ablation, starvation reduced HCRT levels in each cell, without affecting the number of HCRT-positive neurons. Further, starvation did not affect the number of HCRT-positive cells or HCRT levels per cell in surface fish (Fig. 7B). These results indicate that the starvation modulates HCRT levels, rather than the number of cells that produce HCRT. The acute regulation of HCRT by feeding state and lateral-line dependent sensory input demonstrates a unique link between these neuronal systems and those mediating sleep/wake cycles.

**Figure 7.**
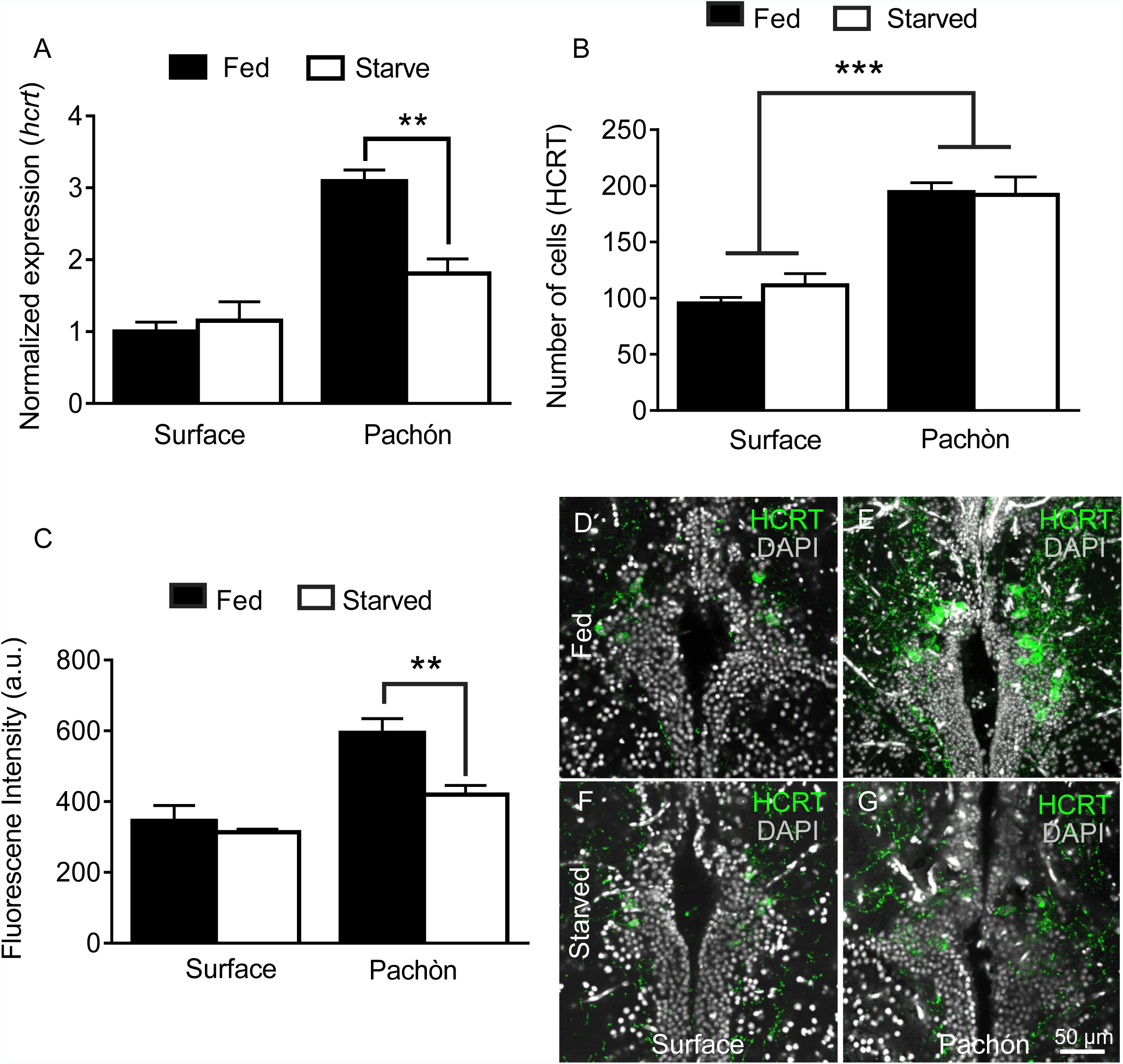
Starvation selectively inhbits HCRT levels in cavefish A. Starvation does not affect *hcrt* expression in surface fish (P>0.832, n=4) while *hcrt* expression is significantly reduced in Pachón cavefish (Pkj >0.001, n=4, 2-way ANOVA, F(1,13)=13.54)) **B.** Fluorescent intensity in HCRT cells was not affected by 30 days starvation in surface fish (P>0.788, n=4). In Pachón cavefish, HCRT neuropeptide was significantly reduced following starvation (P< 0.004, n=4, 2-way ANOVA, F(1,12)=10.17)) **C.** Starvation has no significant effect on total number of HCRT-positive cells in either surface or Pachón cavefish (Surface, P=0.452, n=4; Pachón, P>0.979, n=4, 2-way ANOVA, F(1,11)=3.65)) **D.** Surface control **E.** Pachón control **F.** Surface starved **G.** Pachón Starved. Scale bar =50 µm

## Discussion

Cavefish are a unique model for investigating neural and genetic regulation of sleep, particularly from an evolutionary vantage point. Robust phenotypic differences in sleep have been observed in multiple populations of cavefish, but our findings provide the first evidence for a mechanism that underlies sleep variation between surface and cave fish. Alignment of *hcrt* sequences derived from surface and Pachón cavefish indicate that there are no differences in the protein sequence between the two morphs. Our findings do, however, reveal dramatic differences in *hcrt* expression and neuron number between surface fish and cavefish. These findings suggest that functional differences between surface fish and cavefish likely occur at the level of changes in genomic enhancers or neuronal connectivity, which affect *hcrt* expression. Because our findings also reveal an increased number of HCRT-positive neurons in early development, it is also likely that developmental differences between the brains of surface and cavefish underlie differences in *hcrt* function. Examination of cell body number in 6 dpf fry reveals increased HCRT-positive neurons in cavefish, indicating HCRT differences are present during early development. In agreement with these findings, broad anatomical differences in forebrain structure have previously been documented between surface fish and cavefish including an expanded hypothalamus (Menuet et al., 2007). Therefore, it is likely that developmentally-derived differences in the number of HCRT-positive neurons and modified hypothalamic neural circuitry contribute to sleep loss in cavefish.

Multiple lines of evidence presented here support a robust role for evolved differences in HCRT function in the evolution of sleep loss in Pachón cavefish. Pharamacological inhibition of HCRT signaling, targeted knockdown of HCRT using morpholinos, or genetic inhibition of HCRT neurons promote sleep in Pachón cavefish without significantly affecting sleep in surface fish. While manipulations that inhibit HCRT signaling have potent affects on cavefish sleep, it does not rule out a possible role for HCRT modulation of sleep in surface fish. All three manipulations employed are likely to only partially disrupt HCRTR signaling, and it is possible that complete inhibition of HCRTR signaling would increase sleep in surface fish. Indeed, Hcrt is highly conserved and has been shown to promote wakefulness in species including in animals ranging from zebrafish to mammals (Mieda et al., 2004; Prober et al., 2006). The finding presented here extend studies in mammalian and zebrafish models, and suggest that regulation of HCRT signaling may be subject to evolutionary pressure, and implicate it as a ‘hot-spot’ for variation in sleep throughout the animal kingdom.

While the neural processes regulating HCRT activity are not fully understood, growing evidence suggests these neurons integrate sleep-wake regulation with responses to sensory stimuli (Appelbaum et al., 2007; Mileykovskiy et al., 2005; Woods et al., 2014). In mice, HCRT neurons are transiently activated by sound, feeding, and cage exploration, suggesting that HCRT neurons are generally regulated by external stimuli (Mileykovskiy et al., 2005). Further in zebrafish, HCRT neuron activation is associated with periods of wakefulness (Naumann et al., 2010), overexpression of HCRT enhances locomotor response to diverse sensory stimuli (Prober et al., 2006; Woods et al., 2014), while ablation of HCRT neurons reduces response to a sound stimulus (Elbaz et al., 2012; Naumann et al., 2010; Prober et al., 2006; Woods et al., 2014), suggesting that HCRT neurons link sensory responsiveness to sleep-wake behavior. Our findings reveal that ablation of the lateral line in cavefish reduces *hcrt* transcript and neuropeptide abundance to levels indistinguishable from their surface fish conspecifics, indicating that lateral line input is a potent regulator of *hcrt* production in cavefish. These discoveries suggest that evolution of sensory systems can affect central brain processes that regulate behavior, and provide further support that HCRT neurons integrate sensory stimuli to modulate sleep and arousal.

While a full understanding of the neural circuitry regulating HCRT neurons has not been determined, these neurons project to numerous areas implicated in behavioral regulation, including the periventricular hypothalamus, raphe, and thalamic nuclei (Panula, 2010). Evidence suggests that the wake-promoting role of HCRT neurons is dependent on norepinephrine (Carter et al., 2012; Singh et al., 2015), and optogenetic activation of HCRT neurons activates the locus coeruleus (Singh et al., 2015), raising the possibility that activation of this arousal pathway is enhanced in Pachón cavefish. We previously demonstrated that treating cavefish with the β-adrenergic inhibitor propranolol restores sleep in cavefish without affecting sleep in surface fish (Duboué et al., 2012), similar to findings observed in this study in fish treated with the HCRTR inhibitor TCS0X229. Therefore, it is possible that differences in norepinephrine signaling contribute to sleep loss in cavefish. Further investigation of the synergistic effects of norepinephrine and *hcrt* in surface and cavefish, and the effects of their pre-supposed interaction on feeding- and sensory-mediated *hcrt* production, will be critical in our understanding of how sleep changes can be driven by alterations in the environment.

In addition to its role in sleep-wake regulation, *hcrt* neurons regulate feeding and metabolic function, raising the possibility that HCRT neurons are integrators of sleep and metabolic state. Previous findings reveal that injection of HCRT peptide increases food consumption in cavefish, suggesting the consummatory behavior induced by HCRT in mammals is conserved in *A. mexicanus* (Penney and Volkoff, 2014; Wall and Volkoff, 2013). In addition, studies in mammals and zebrafish suggest HCRT neurons are regulated by the adipose peptide hormone, Leptin (Leinninger et al., 2011; Levitas-Djerbi et al., 2015). Adipose levels in cavefish are elevated compared to surface fish (Aspiras et al., 2015), and it is possible that prolonged starvation reduces leptin levels, thereby inhibiting *hcrt* expression to promote sleep in cavefish. These findings suggest that cavefish may provide a useful model for examining the leptin-Hcrt axis and, more generally, interactions between sleep and metabolic function.

Our findings specifically examine neuronal mechanisms underlying sleep loss in the Pachón cave populations. Both morphological and genomic data suggest Pachón cavefish are one of the oldest, and most troglomorphic of the 29 *A. mexicanus* cavefish populations (Bradic et al., 2012; Dowling et al., 2002; Ornelas-García et al., 2008; Strecker et al., 2003). We have also demonstrated evolutionary convergence on sleep loss in other populations of cavefish, including Molino, Tinaja and Chica cave populations (Jaggard et al., 2017). However, ablation of the lateral line has no effect on sleep in Molino, Tinaja and Chica populations, suggesting distinct neural mechanism underlie sleep loss between Pachón cavefish, and other cavefish populations assayed (Jaggard et al., 2017). Future studies will reveal if enhanced HCRT function represents a conserved mechanism for sleep loss, or that sleep loss in other fish populations is HCRT independent.

## Materials and Methods

### Fish maintenance and rearing

Animal husbandry was carried out as previously described (Borowsky, 2008b) and all protocols were approved by the IACUC Florida Atlantic University. Fish were housed in the Florida Atlantic University core facilities at 21°C ± 1°C water temperature throughout rearing for behavior experiments (Borowsky, 2008b). Lights were kept on a 14:10 hr light-dark cycle that remained constant throughout the animal’s lifetime. Light intensity was kept between 25-40 Lux for both rearing and behavior experiments. All fish used for experiments were raised to adulthood and housed in standard 18-37 L tanks. Adult fish were fed a mixture diet of black worms to satiation twice daily at zeitgeber time (ZT) 2 and ZT12, (Aquatic Foods, Fresno, CA,) and standard flake fish food during periods when fish were not being used for behavior experiments or breeding (Tetramine Pro).

### Sleep behavior

Adult fish were recorded in standard conditions in 10 L tanks with custom-designed partitions that allowed for five fish (2 L/fish) to be individually housed in each tank as previously described (Yoshizawa et al., 2015). Recording chambers were illuminated with custom-designed IR LED source (Infrared 850 nm 5050 LED Strip Light, Environmental Lights). After a 4-5 day acclimation period, behavior was recorded for 24 h beginning at ZT0-ZT2. Videos were recorded at 15 frames/s using a USB webcam (LifeCam Studio 1080p HD Webcam, Microsoft) fitted with a zoom lens (Zoom 7000, Navitar). An IR high-pass filter (Edmund Optics Worldwide) was placed between the camera and the lens to block visible light. For larval fish recordings, individual fish were placed in 12 well tissue culture plates (BD Biosciences). Recording chambers were lit with a custom-designed IR LED light strip placed beneath the recording platform. Larvae were allowed to acclimate for 24 hours before starting behavioral recordings. Videos were recorded using Virtualdub, a video-capturing software (Version 1.10.4) and were subsequently processed using Ethovision XT 9.0 (Noldus, IT). Water temperature and chemistry were monitored throughout recordings, and maintained at standard conditions in all cases. Ethovision tracking was setup as previously described (Yoshizawa et al., 2015). Data was then processed using Perl scripts (v5.22.0, developed on-site) and Excel macro (Microsoft) (Yoshizawa et al., 2015). These data were used to calculate sleep information by finding bouts of immobility of 60 s and greater, which are correlated with increased arousal threshold, one of the hallmarks of sleep (Yoshizawa et al., 2015). For drug treatment studies, fish were allowed normal acclimation periods, followed by 24 h of baseline recording. At ZT0 fish were treated with either control dimethyl sulfoxide vehicle (0.1% DMSO) or freshly prepared TCS0X229 (Tocris), EMPA (Tocris), Suvorexant (Adooq), or YNT-185 (Adooq) (Hoyer et al., 2013; Kummangal et al., 2013; Malherbe et al., 2009; Plaza-Zabala et al., 2012) diluted to a final concentration of 1- 30 µM into each recording chamber and behavior was recorded for 24 h.

### Injection procedures

The *hcrt* morpholino sequence 5’-TGGGCTTGGTGTGATCACCTGTCAT-3’ was designed (Gene Tools, LLC) based on the available sequence ENSAMXG00000000473 (Ensembl). The HCRT-MO targets the first 25-bp of the *hcrt* open reading frame (ORF) to block translation via steric hindrance. Control injections were performed using a standard scrambled sequence 5’CCTCTTACCTCAGTTACAATTTATA-3’ (Gene Tools, LLC). Morpholino injections were carried out using 2 ng of MO in a 1 nl volume, in accordance with previously published methods (Bilandzija et al., 2013; Gross et al., 2009). Surface and Pachón cavefish embryos were injected at 1-2 cell stage using a pulled borosilicate capillary with a Warner PLI-A100 picoinjector. Survival of all embryos was monitored every 6 h for the first 96 h of development until behavior was recorded over the next 24 h. *hcrt:Gal4; uas:BoTxBLC-GFP* injections were carried out as follows: Separate plasmids containing the Gal4 and the BoTx transgenes were coinjected into 1-4 cell stage embryos using a pulled borosilicate capillary with Warner PLI-A100 picoinjector at concentration of 25 ng/µL. *Tol2* mRNA was coinjected in the cocktail at a concentration of 25 ng/µL. *uas:BoTx-BLC-GFP* was injected alone at 25 ng/µL as a negative control. All injected fish were raised to 25 dpf in standard conditions, when behavioral recordings were carried out. Brains from all fish recorded for behavior were dissected and processed for immunohistochemistry in order to quantify by GFP the number of cells expressing *hcrt:Gal4; uas:BoTx-BLC-GFP*.

### Vital dye labeling and lateral line ablation

Fish were treated with 0.002% gentamicin sulfate (Sigma Aldrich 1405-41-0) as previously described (Van Trump et al., 2010). Following baseline sleep recording and neuromast imaging, fish were bathed in gentamicin for 24 h. Following the treatment, a complete water change was administered and behavior was again recorded for 24 h. Fish treated with gentamicin were housed in separate tanks for at least 1 month after treatment in order to avoid contamination. Lateral line re-growth was measured with DASPEI staining two weeks following ablation to confirm that there were no long-term effects from the ablation treatments.

### Sequence analysis

To compare Hcrt sequences in different species, we aligned the Hcrt protein sequences of *A. mexicanus* surface fish <u>(SRR639083.116136.2, SRA)</u> and Pachón cavefish (ENSAMXP00000000478) to that of zebrafish (ENSDARP00000095322), Medaka (ENSORLP00000004866), Tetraodon (ENSTNIP00000014660), mouse (ENSMUSP00000057578) and human (ENSP00000293330). Protein alignment, neighbor joining tree (cladogram) and sequences analyses were performed with Clustal Omega (v.1.2.1, EMBL-EBI, (Sievers et al., 2011)). HCRT domains (PF02072/IPR001704) were determined using Ensembl genome browser (v.83, EMBL-EBI/Sanger) and PFam/Interpro (v.28.0, EMBL-EBI).

### Quantitative PCR (qPCR)

To measure levels of *hcrt* mRNA, whole brains of one-two year old fish were extracted immediately after behavior was recorded (ZT2). After extraction, individual brains were frozen and homogenized in trizol (QIAGEN, Valencia,CA). RNA was extracted with an RNeasy Mini Kit (QIAGEN, Valencia,CA). All RNA samples were standardized to 10 ng/µL and cDNA synthesis was carried out using iScript (BioRad, Redmond, WA). RT-qPCR was carried out using SsoAdvanced Universal SYBR Green Supermix (BioRad, Redmond, WA). qPCR primers were: *hcrt* forward 5’-CAT-CTC-CTC-AGC-CAA-GGT-TT-3’, *hcrt* reverse 5’-TAG-AGT-CCG-TGC-TGT-TAC-ACC-3’. Two housekeeping genes were amplified with the following primers: *rpl13α* forward 5’-TCT-GGA-GGA-CTG-TAA-GAG-GTA-TGC-3’, *rpl13α* reverse 5’-AGA-CGC-ACA- ATC-TTG-AGA-GCA-G-3’; *gapdh* forward 5’-GTG-TCC-ACA-GAC-TTC-AAC-GG-3’, *gapdh* reverse 5’CAT-TGT-CAT-ACC-ATG-TCA-CCA-G-3’. The following qPCR protocol was run on a Bio Rad CFX96 with a C1000 thermal cycler: 95.0°C for 3 min followed by a plate read at 95.0°C for 10 s to 53.3°C for 30 s followed by a plate read repeated 39 times. All samples were compiled into Bio Rad CFX manager gene study (version 3.1) to account for inter-run calibration. All samples were normalized to one (relative to surface fish controls) Housekeeping genes were validated using Bio Rad CFX manager and found M values of 0.82 for *rpl13α,* and 0.57 for *gapdh*, falling well within the acceptable range for quality reference genes (Vandesompele et al., 2002).

### Immunohistochemistry

Following euthanasia in MS-222 (Sigma) and ice-water, brains were immediately dissected from adults in ice-cold PBS and fixed overnight in 4% paraformaldehyde/1x PBS (PFA). Adult brains were then placed in 20% sucrose for cryoprotection overnight or until the brains sunk to the bottom of the well (Kaslin et al, 2004). Whole brains were then flash frozen and mounted in OCT compound (23-730-571 Fisher scientific) for sectioning. Whole brains were serial sectioned in 50 µm slices, all slices were floated in PBS to rinse out embedding solution. Slices were then washed in 0.5% Triton-X 100/PBS (PBT) for 3X 15 minutes and co-incubated in 0.5% PBT and 2% bovine serum albumin (BSA) (Sigma) with primary antibody anti-ORX-A 1:2000 (AB3704 EMD Millipore) overnight at 4°C with gentle shaking. The slices were rinsed again in 0.5% PBT for 3X 15 minutes and placed in secondary antibody (1:600 goat anti-rabbit 488; Life Technologies) for 90 minutes at room temperature. Slices were mounted on slides in Vectashield with DAPI (VectorLabs) and imaged on a Nikon A1 confocal microscope. Whole-mount larvae were fixed overnight in 4% PFA, rinsed 3X 15 min in 0.5% PBT, then placed in 0.5% PBT with 2% BSA and primary antibody anti-ORX-A 1:2000 overnight at 4°C. Following 3X 15 min 0.5% PBT rinse, larvae were incubated in secondary antibody at 1:600. Larvae were then placed in Vectashield with DAPI until mounted in 2% low melt agarose (Sigma) for imaging. All samples were imaged in 2 µm sections and are presented as the Z-stack projection through the entire brain. For quantification of HCRT levels, all hypothalamic slices were imaged in 2 µm sections, merged into a single Z-stack using maximum intensity projections (Nikon Elements), and the total brain fluorescence was determined by creating individual ROIs for all soma expressing HCRT, background intensity was subtracted in order correct for non-specific fluorescence. All imaging analysis was carried out using Nikon Elements (v. 4.50).

### Statistics

Two-way ANOVA tests were carried out to test the effects of pharmacological and starvation paradigms among different groups and populations on behavior. Each was modeled as a function of genotype (Surface and Pachón) and genotype by treatment interaction (TCS, gentamicin, or starvation, respectively). Significance for all tests was set at p< 0.05. When the ANOVA test detected significance, the Holm-Sidak multiple comparison post-test was carried out to correct for the number of comparisons. For comparison of two baseline groups, non-parametric t-tests were carried out to test for significance. Each experiment was repeated independently at least three times. All replicates were biological replicates run independently from one another. No data was excluded, and no statistical outliers were removed. All statistical analyses were carried out using SPSS (IBM, 22.0) or InStat software (GraphPad 6.0). Power analyses were performed to ensure that we had sufficient N to detect significant differences at a minimum of 80% power at the 0.05 threshold using Graphpad InStat.

## Acknowledgements

This work was funded by National Science Foundation Award IOS-125762 to ACK. The authors are grateful to Masato Yoshizawa (Hawai’i) and Lior Appelbaum (Bar Illan University) for technical guidance and valuable discussion. The Department of Comparative Medicine at FAU for support maintaining the fish facility. The authors received reagents from Kochi Kawakami (Sokendai) and Lior Appelbaum (Bar Illan University).

## Supplementary Figures

**Figure 1, Figure Supplement 1. HCRT sequence is identical between surface fish and Pachón cavefish. A.** Sequence alignment for HCRT in *A. mexicanus* reveals a homology of 35- 48% percent identity compared to other fish species including zebrafish and medaka, and a 35% percent identity conserved compared to mammalian HCRT. **B.** Housekeeping genes used for all q-PCR experiments are not significantly altered between Surface and Cavefish. Expression was normalized to Surface fish.

**Figure 1, Figure Supplement 2. HCRT peptide levels are increased in Pachón cavefish. A.** Surface fish whole brain from sagittal 2 µm confocal slice **B.** Pachón cavefish whole brain sagittal 2 µm confocal slice. **C.** Surface fish hypothamus sagittal 2 µm confocal slice. **D.** Pachón cavefish hypothalmus 2 µm confocal slice

**Figure 1, Figure Supplement 3. Hypocretin levels are increased in early development in Pachón cavefish. A.** Five days post fertilization whole-mount surface fish immunostained with anti-HCRT (green) and DAPI (white). **B.** Whole mount four dpf Pachón cavefish immunostained with anti-HCRT (Green) and DAPI (white). **C.** HCRT neuropeptide levels are significantly increased in five dpf Pachón larvae compared to surface (Unpaired t-test, t= 3.17, df=10, P< 0.01).

**Figure 3, Figure Supplement 1. HCRT agonist partially alters sleep behavior in juvenile surface fish, but not Pachón cavefish. A.** Total sleep was significantly reduced in surface fish (P=0.001, n=26) while there was no effect with YNT-185 treatment in Pachón cavefish (P=0.997, n=16; 2-way ANOVA, F(1,77)=25.81, P< 0001). **B.** Waking activity was not significantly altered between Surface fish (P=0.228, n=26) or Pachon cavefish (P=0.716, n=16) or with treatment of YNT-185 among either treatment groups (2-way ANOVA, F(1,77)=0.133, P=0.715). **C.** Total sleep bouts were significantly reduced in surface fish with YNT-185 treatment (P=0.0005, n=26). However, there was no alteration in sleep bouts in Pachón cavefish (P=0.989, n=16; 2-way ANOVA, F(1,77)=14.78, P=0.0002). **D.** Average Sleep bout duration was not significantly changed in either Surface fish (P=0.391, n=26) or in Pachón cavefish (P=0.820, n=16; 2-way ANOVA, F(1,77)=2.06, P=0.155).

**Figure 2, Figure Supplement 2 HCRTR2 blockade selectively increases sleep in Pachón cavefish. A** Bright field images of 25 dpf Surface and Pachón cavefish (scale bar represents 200 µm). **B.** TCS treatment did not affect sleep duration in surface fish (1 uM, p=0.782, n=18, 10 uM P>0.891, n=18) while Pachón cavefish increased total sleep duration in response to 10uM TCS treatment (1uM, P>0.116, n=17; 10uM, P< 0.031, n=18, 2-way ANOVA, F(2,88)=12.178). **C-D.** Twenty-four hour sleep profiles of surface fish **(C)** and Pachón cavefish **(D).**

**Figure 5: Supplemental Figure 1: Morpholino injections do not affect survival in development. A.** No significant difference in Surface fish survival from embryo injection to 96 hours post fertilization (Kruskall-Wallis, P=0.719, Dunn’s post test, Wild Type vs. Scramble MO, P=0.862, Scramble MO vs. HCRT MO, P=0.701). **B.** Pachón fish survival from embryo injection compared to controls up to 96 hours post fertilization (Kruskall-Wallis, P=0.429, Dunn’s post test, Wild Type vs. Scramble MO, P=0.299, Scramble MO vs. HCRT MO, P=0.206).

**Figure 6: Supplemental Figure 1: Genetic silencing of subsets of HCRT alters sleep architecture in Pachón cavefish. A.** Total sleep bouts were not significantly altered in Surface fish when HCRT cells were silenced (Wild Type*UAS-BoTx-BLC-GFP, P=0.561, n=23; UAS- BoTx-BLC-GFP* hcrt:GAL4-UAS-BoTx-BLC-GFP, P=0.636, n=14). When HCRT cell were silenced in Pachón cavefish there was a significant increased their total number of sleep bouts (Wild Type*UAS-BoTx-BLC-GFP, P=0.561, n=21; UAS-BoTx-BLC-GFP* hcrt:GAL4-UAS-BoTx-BLC-GFP, P=0.031, n=15; 2-way ANOVA, F(1,91)=32.43, P< 0.0001). **B.** Sleep bout duration was not changed in either Surface fish or Pachon cavefish when HCRT neurons were silenced (Surface Wild Type*UAS-BoTx-BLC-GFP, P>0.999, n=21; Surface UAS-BoTx-BLC-GFP* hcrt:GAL4-UAS-BoTx-BLC-GFP, P=0.743, n=15; Pachón Wild Type*UAS-BoTx-BLC-GFP, P=0.401, n=21; Pachón UAS-BoTx-BLC-GFP* hcrt:GAL4-UAS-BoTx-BLC-GFP, P>0.999, n=15;2-way ANOVA, F(1,91)=3.886, P=0.052).

## References

Adamantidis, A., and de Lecea, L. (2008). Sleep and metabolism: shared circuits, new connections. Trends Endocrinol. Metab. 19, 362–370.

Allada, R., and Siegel, J.M. (2008). Unearthing the phylogenetic roots of sleep. Curr. Biol. 18, R670–R679.

Appelbaum, L., Skariah, G., Mourrain, P., and Mignot, E. (2007). Comparative expression of p2x receptors and ecto-nucleoside triphosphate diphosphohydrolase 3 in hypocretin and sensory neurons in zebrafish. Brain Res. 1174, 66–75.

Asakawa, K., and Kawakami, K. (2008). Targeted gene expression by the Gal4-UAS system in zebrafish. Dev. Growth Differ. 50, 391–399.

Aspiras, A., Rohner, N., Marineau, B., Borowsky, R., and Tabin, J. (2015). Melanocortin 4 receptor mutations contribute to the adaptation of cavefish to nutrient-poor conditions. Proc. Natl. Acad. Sci. 112, 9688–73.

Auer, T.O., Xiao, T., Bercier, V., Gebhardt, C., Duroure, K., Concordet, J.P., Wyart, C., Suster, M., Kawakami, K., Wittbrodt, J., et al (2015). Deletion of a kinesin I motor unmasks a mechanism of homeostatic branching control by neurotrophin-3. Elife 4.

Betschart, C., Hintermann, S., Behnke, D., Cotesta, S., Fendt, M., Gee, C.E., Jacobson, L.H., Laue, G., Ofner, S., Chaudhari, V., et al (2013). Identification of a Novel Series of Orexin Receptor Antagonists with a Distinct Effect on Sleep Architecture for the Treatment of Insomnia. J Med Chem 56, 7590–7609.

Bilandzija, H., Ma, L., Parkhurst, A., and Jeffery, W. (2013). A potential benefit of albinism in Astyanax cavefish: downregulation of the oca2 gene increases tyrosine and catecholamine levels as an alternative to melanin synthesis. PLoS One 8, e80823.

Bill, B.R., Petzold, A.M., Clark, K.J., Schimmenti, L. a, and Ekker, S.C. (2009). A primer for morpholino use in zebrafish. Zebrafish 6, 69–77.

Borowsky, R. (2008a). Restoring sight in blind cavefish. Curr. Biol. 18, R23–R24.

Borowsky, R. (2008b). Handling Astyanax mexicanus eggs and fry. Cold Spring Harb. Protoc. 3.

Bradic, M., Beerli, P., Garcia-de Leon, F.J., Esquivel-Bobadilla, S., and Borowsky, R.L. (2012). Gene flow and population structure in the Mexican blind cavefish complex (*Astyanax mexicanus*). BMC Evol. Biol. 12, 9 doi:10.1186/1471-2148-12-9

Brand, A.H., and Perrimon, N. (1993). Targeted gene expression as a means of altering cell fates and generating dominant phenotypes. Development 118, 401–415.

Brunger, a T., Jin, R., and Breidenbach, M. a (2008). Highly specific interactions between botulinum neurotoxins and synaptic vesicle proteins. Cell. Mol. Life Sci. 65, 2296–2306.

Campbell, S.S., and Tobler, I. (1984). Animal sleep: a review of sleep duration across phylogeny. Neurosci. Biobehav. Rev. 8, 269–300.

Capellini, I., Barton, R.A., McNamara, P., Preston, B.T., and Nunn, C.L. (2008). Phylogenetic analysis of the ecology and evolution of mammalian sleep. Evolution (N. Y). 62, 1764–1776.

Carter, M.E., Brill, J., Bonnavion, P., Huguenard, J.R., Huerta, R., and de Lecea, L. (2012). Mechanism for Hypocretin-mediated sleep-to-wake transitions. Proc. Natl. Acad. Sci. U. S. A. 109, E2635–44.

Chemelli, R.M., Willie, J.T., and Sinton, C.M. et al (1999). Narcolepsy in orexin knockout mice: Molecular genetics of sleep regulation. Cell 98, 437–451.

Chen, S., Chiu, C., McArthur, K., Fetcho, J., and Prober, D. (2016). TRP channel mediated neuronal activation and ablation in freely behaving zebrafish. Nat Methods 2, 147–150.

Dowling, T.E., Martasian, D.P., and Jeffery, W.R. (2002). Evidence for multiple genetic forms with similar eyeless phenotypes in the blind cavefish, Astyanax mexicanus. Mol. Biol. Evol. 19, 446–455.

Duboué, E.R., Keene, A.C., and Borowsky, R.L. (2011). Evolutionary convergence on sleep loss in cavefish populations. Curr. Biol. 21, 671–676.

Duboué, E.R., Borowsky, R.L., and Keene, A.C. (2012). β-adrenergic signaling regulates evolutionarily derived sleep loss in the mexican cavefish. Brain. Behav. Evol. 80, 233–243.

Elbaz, I., Yelin-Bekerman, L., Nicenboim, J., Vatine, G., and Appelbaum, L. (2012). Genetic Ablation of Hypocretin Neurons Alters Behavioral State Transitions in Zebrafish. J. Neurosci. 32, 12961–12972.

Gross, J.B. (2012). The complex origin of *Astyanax* cavefish. BMC Evol. Biol. 12, 105 doi:10.1186/1471-2148-12-105.

Gross, J.B., and Wilkens, H. (2013). Albinism in phylogenetically and geographically distinct populations of *Astyanax* cavefish arises through the same loss-of-function Oca2 allele. Heredity (Edinb). 111, 122–130.

Gross, J.B., Borowsky, R., and Tabin, C.J. (2009). A Novel Role for *Mc1r* in the Parallel Evolution of Depigmentation in Independent Populations of the Cavefish *Astyanax mexicanus*. PLoS Genet 5, e1000326.

Gross, J.B., Furterer, A., Carlson, B.M., and Stahl, B.A. (2013). An Integrated Transcriptome-Wide Analysis of Cave and Surface Dwelling Astyanax mexicanus. PLoS One 8.

Hartmann, E.L. (1973). The Functions of Sleep (Yale University Press).

Hoyer, D., Dürst, T., Fendt, M., Jacobson, L.H., Betschart, C., Hintermann, S., Behnke, D., Cotesta, S., Laue, G., Ofner, S., et al (2013). Distinct effects of IPSU and suvorexant on mouse sleep architecture. Front. Neurosci.

Irukayama-Tomobe, Y., Ogawa, Y., Tominaga, H., Ishikawa, Y., Hosokawa, N., Ambai, S., Kawabe, Y., Uchida, S., Nakajima, R., Saitoh, T., et al (2017). Nonpeptide orexin type-2 receptor agonist ameliorates narcolepsy-cataplexy symptoms in mouse models. Proc. Natl. Acad. Sci. 114, 201700499.

Jaggard, J., Robinson, B., Stahl, B., Oh, I., Masek, P., Yoshizawa, M., and Keene, A. (2017). The lateral line confers evolutionarily derived sleep loss in the Mexican cavefish. J. Exp. Biol. in press.

Jeffery, W.R. (2009). Regressive Evolution in *Astyanax* Cavefish. Annu. Rev. Genet. 43, 25–47.

Kawakami, K., Shima, A., and Kawakami, N. (2000). Identification of a functional transposase of the Tol2 element, an Ac-like element from the Japanese medaka fish, and its transposition in the zebrafish germ lineage. Proc. Natl. Acad. Sci. 97, 11403–11408.

Keene, A., Yoshizawa, M., and McGaugh, S. (2015). Biology and Evolution of the Mexican Cavefish (New York: Academic Press).

Kowalko, J.E., Rohner, N., Rompani, S.B., Peterson, B.K., Linden, T.A., Yoshizawa, M., Kay, E.H., Weber, J., Hoekstra, H.E., Jeffery, W.R., et al (2013). Loss of schooling behavior in cavefish through sight-dependent and sight-independent mechanisms. Curr. Biol. 23, 1874–1883.

Kulpa, M., Bak-Coleman, J., and Coombs, S. (2015). The lateral line is necessary for blind cavefish rheotaxis in non-uniform flow. J. Exp. Biol. 218, 1603–1612.

Kummangal, B.A., Kumar, D., and Mallick, H.N. (2013). Intracerebroventricular injection of orexin-2 receptor antagonist promotes REM sleep. Behav. Brain Res. 237, 59–62.

Leinninger, G.M., Opland, D.M., Jo, Y.H., Faouzi, M., Christensen, L., Cappellucci, L.A., Rhodes, C.J., Gnegy, M.E., Becker, J.B., Pothos, E.N., et al (2011). Leptin action via neurotensin neurons controls orexin, the mesolimbic dopamine system and energy balance. Cell Metab. 14, 313–323.

Levitas-Djerbi, T., Yelin-Bekerman, L., Lerer-Goldshtein, T., and Appelbaum, L. (2015). Hypothalamic leptin-neurotensin-hypocretin neuronal networks in zebrafish. J. Comp. Neurol. 523, 831–848.

Lin, L., Faraco, J., Li, R., Kadotani, H., Rogers, W., Lin, X., Qiu, X., De Jong, P.J., Nishino, S., and Mignot, E. (1999). The sleep disorder canine narcolepsy is caused by a mutation in the hypocretin (orexin) receptor 2 gene. Cell 98, 365–376.

Malherbe, P., Borroni, E., Gobbi, L., Knust, H., Nettekoven, M., Pinard, E., Roche, O., Rogers-Evans, M., Wettstein, J.G., and Moreau, J.L. (2009). Biochemical and behavioural characterization of EMPA, a novel high-affinity, selective antagonist for the OX 2 receptor. Br. J. Pharmacol. 156, 1326–1341.

McGaugh, S., Gross, J.B., Aken, B., Blin, M., Borowsky, R., Chalopin, D., Hinaux, H., Jeffery, W., Keene, A., Ma, L., et al (2014). The cavefish genome reveals candidate genes for eye loss. Nat. Commun. 5, 5307.

Menuet, A., Alunni, A., Joly, J.-S., Jeffery, W.R., and Rétaux, S. (2007). Expanded expression of Sonic Hedgehog in Astyanax cavefish: multiple consequences on forebrain development and evolution. Development 134, 845–855.

Mieda, M., Willie, J.T., Hara, J., Sinton, C.M., Sakurai, T., and Yanagisawa, M. (2004). Orexin peptides prevent cataplexy and improve wakefulness in an orexin neuron-ablated model of narcolepsy in mice. Proc. Natl. Acad. Sci. 101, 4649–4654.

Mileykovskiy, B.Y., Kiyashchenko, L.I., and Siegel, J.M. (2005). Behavioral correlates of activity in identified hypocretin/orexin neurons. Neuron 46, 787–798.

Mitchell, R.W., Russell, W.H., and Elliott, W.R. (1977). Mexican eyeless characin fishes, genus Astyanax: Environment, distribution, and evolution. (Texas: Texas Tech Press).

Mochizuki, T., Arrigoni, E., Marcus, J.N., Clark, E.L., Yamamoto, M., Honer, M., Borroni, E., Lowell, B.B., Elmquist, J.K., and Scammell, T.E. (2011). Orexin receptor 2 expression in the posterior hypothalamus rescues sleepiness in narcoleptic mice. Proc. Natl. Acad. Sci. U. S. A. 108, 4471–4476.

Musiek, E.S., Xiong, D.D., and Holtzman, D.M. (2015). Sleep, circadian rhythms, and the pathogenesis of Alzheimer disease. Exp. Mol. Med. 47, e148.

Nagahara, T., Saitoh, T., Kutsumura, N., Irukayama-Tomobe, Y., Ogawa, Y., Kuroda, D., Gouda, H., Kumagai, H., Fujii, H., Yanagisawa, M., et al (2015). Design and Synthesis of Non-Peptide, Selective Orexin Receptor 2 Agonists. J. Med. Chem. 58, 7931–7937.

Naumann, E.A., Kampff, A.R., Prober, D.A., Schier, A.F., and Engert, F. (2010). Monitoring neural activity with bioluminescence during natural behavior. Nat. Neurosci. 13, 513–520.

Ornelas-García, C.P., Domínguez-Domínguez, O., and Doadrio, I. (2008). Evolutionary history of the fish genus Astyanax Baird & Girard (1854) (Actinopterygii, Characidae) in Mesoamerica reveals multiple morphological homoplasies. BMC Evol. Biol. 8, 340.

Panula, P. (2010). Hypocretin/orexin in fish physiology with emphasis on zebrafish. In Acta Physiologica, pp. 381–386.

Penney, C., and Volkoff, H. (2014). Peripheral injections of cholecystokinin, apelin, ghrelin and orexin in cavefish (Astyanax fasciatus mexicanus): effects on feeding and on the brain expression levels of tyrosine hydroxylase, mechanistic target of rapamycin and appetite-related hormones. Gen Comp Endocrinol 196, 34–40.

Plaza-Zabala, A., Flores, Á., and Maldonado, R. et al (2012). Hypocretin/orexin signaling in the hypothalamic paraventricular nucleus is essential for the expression of nicotine withdrawal. Biol. Psychiatry 71, 214–223.

Prober, D.A., Rihel, J., Onah, A.A., Sung, R.-J., and Schier, A.F. (2006). Hypocretin/orexin overexpression induces an insomnia-like phenotype in zebrafish. J. Neurosci. 26, 13400–13410.

Protas, M.E., Hersey, C., Kochanek, D., Zhou, Y., Wilkens, H., Jeffery, W.R., Zon, L.I., Borowsky, R., and Tabin, C.J. (2006). Genetic analysis of cavefish reveals molecular convergence in the evolution of albinism. Nat. Genet. 38, 107–111.

Rihel, J., Prober, D.A., Arvanites, A., Lam, K., Zimmerman, S., Jang, S., Haggarty, S.J., Kokel, D., Rubin, L.L., Peterson, R.T., et al (2010). Zebrafish behavioral profiling links drugs to biological targets and rest/wake regulation. Science 327, 348–351.

Scheer, N., and Campos-Ortega, J.A. (1999). Use of the Gal4-UAS technique for targeted gene expression in the zebrafish. Mech. Dev. 80, 153–158.

Siegel, J.M. (2005). Clues to the functions of mammalian sleep. Nature 437, 1264–1271.

Sievers, F., Wilm, A., Dineen, D., Gibson, T.J., Karplus, K., Li, W., Lopez, R., McWilliam, H., Remmert, M., Söding, J., et al (2011). Fast, scalable generation of high-quality protein multiple sequence alignments using Clustal Omega. Mol. Syst. Biol. 7, 539.

Singh, C., Oikonomou, G., and Prober, D. (2015). Norepinephrine is required to promote wakefulness and for hypocretin-induced arousal in zebrafish. Elife 4, e070000.

Strecker, U., Bernatchez, L., and Wilkens, H. (2003). Genetic divergence between cave and surface populations of Astyanax in Mexico (Characidae, Teleostei). Mol. Ecol. 12, 699–710.

Van Trump, W.J., Coombs, S., Duncan, K., and McHenry, M.J. (2010). Gentamicin is ototoxic to all hair cells in the fish lateral line system. Hear. Res. 261, 42–50.

Tsujino, N., and Sakurai, T. (2013). Role of orexin in modulating arousal, feeding, and motivation. Front. Behav. Neurosci. 7, 28.

Vandesompele, J., De Preter, K., Pattyn, ilip, Poppe, B., Van Roy, N., De Paepe, A., and Speleman, rank (2002). Accurate normalization of real-time quantitative RT-PCR data by geometric averaging of multiple internal control genes. Genome Biol. 3, 34–1.

Wall, A., and Volkoff, H. (2013). Effects of fasting and feeding on the brain mRNA expressions of orexin, tyrosine hydroxylase (TH), PYY and CCK in the Mexican blind cavefish (Astyanax fasciatus mexicanus). Gen. Comp. Endocrinol. 183, 44–52.

Wong, K.K.Y., Ng, S.Y.L., Lee, L.T.O., Ng, H.K.H., and Chow, B.K.C. (2011). Orexins and their receptors from fish to mammals: A comparative approach. Gen. Comp. Endocrinol. 171, 124–130.

Woods, I.G., Schoppik, D., Shi, V.J., Zimmerman, S., Coleman, H.A., Greenwood, J., Soucy, E.R., and Schier, A.F. (2014). Neuropeptidergic signaling partitions arousal behaviors in zebrafish. J. Neurosci. 34, 3142–3160.

Yokobori, E., Kojima, K., Azuma, M., Kang, K.S., Maejima, S., Uchiyama, M., and Matsuda, K. (2011). Stimulatory effect of intracerebroventricular administration of orexin A on food intake in the zebrafish, Danio rerio. Peptides 32, 1357–1362.

Yokogawa, T., Marin, W., Faraco, J., Pézeron, G., Appelbaum, L., Zhang, J., Rosa, F., Mourrain, P., and Mignot, E. (2007). Characterization of sleep in zebrafish and insomnia in hypocretin receptor mutants. PLoS Biol. 5, 2379–2397.

Yoshizawa, M. (2015). Behaviors of cavefish offer insight into developmental evolution. Mol. Reprod. Dev. 82, 268–280.

Yoshizawa, M., Gorički, Š., Soares, D., and Jeffery, W.R. (2010). Evolution of a behavioral shift mediated by superficial neuromasts helps cavefish find food in darkness. Curr. Biol. 20, 1631–1636.

Yoshizawa, M., Robinson, B., Duboue, E., Masek, P., Jaggard, J., O’Quin, K., Borowsky, R., Jeffery, W., and Keene, A. (2015). Distinct genetic architecture underlies the emergence of sleep loss and prey-seeking behavior in the Mexican cavefish. BMC Biol. 20, 15.

Borowsky, R. (2008a). Restoring sight in blind cavefish. Curr. Biol. 18, R23–R24.

Borowsky, R. (2008b). Handling Astyanax mexicanus eggs and fry. Cold Spring Harb. Protoc. 3.

Hartmann, E.L. (1973). The Functions of Sleep (Yale University Press).

Jeffery, W.R. (2009). Regressive Evolution in *Astyanax* Cavefish. Annu. Rev. Genet. 43, 25–47.

Siegel, J.M. (2005). Clues to the functions of mammalian sleep. Nature 437, 1264–1271.

